# Structure and Function of Lens Suture Examined by 2-photon Fluorescence Microscopic Imaging

**DOI:** 10.1101/2025.09.08.675023

**Authors:** Qinrong Zhang, Jun Zhu, Taishi Painter, Chun-Hong Xia, Na Ji, Xiaohua Gong

## Abstract

We have applied 2-photon fluorescence microscopy (2PFM) to investigate the structures and functions of mouse eye lenses, especially lens sutures, using wildtype (WT) and KLPH-KO suture-cataract lenses *in vivo*. Dynamic structures of lens sutures have been hypothesized to act as pathways carrying fluid containing ions, nutrients, and other factors as part of microcirculation for maintaining lens homeostasis and transparency. 3D imaging results show both typical “Y” and “double Y” shaped suture structures in WT lenses, diverse suture patterns including “star” shapes with variations appearing at different depths, and misaligned suture planes in KLPH-KO lenses. The variability in suture patterns, quantified using the mean of image stacks’ similarity indices (SSIM) to the mean projection of the stack, reveals a significant increase (p<0.05) in pattern randomization for KLPH-KO lenses. Both WT and KLPH-KO lenses exhibit voids around sutures and enlarged vacuoles throughout the lens structure *in vivo*. Pathological features like irregular and larger central voids containing 2~5-µm-diameter amorphous structures that might result from membrane protein remnants/aggregates in the KLPH-KO mouse eye lens were observed. In conclusion, this work provides new morphological markers for characterizing suture cataract and fiber pathological features. Lens sutures are stabilized structures that are impermeable to dyes, and the extracellular spaces of lens peripheral fibers provide efficient diffusion pathways for lens microcirculation.

## Introduction

Normal vision relies on the ocular lens, which is a transparent, biconvex structure with appropriate refractive indices located behind the iris, primarily functioning to transmit a focused light image onto the retina for subsequent visual perception^1^. The lens is composed of a bulk of elongated fibers covered with an epithelial cell monolayer at the anterior hemisphere, wrapped with a basement membrane, named the capsule, on its surface. The cellular organization and optical properties of lens fiber cells determine the lens’s optical properties. Mammalian lenses like mouse and human lenses grow bigger throughout their lifetime. Lens growth relies on epithelial cells differentiating into elongating fiber cells precisely overlaid on top of previous elongated fibers at the lens equator. The ends of newly elongating fiber cells extend anteriorly and posteriorly along the capsule towards the polar region^2–5^. Fully elongated fiber ends eventually detach from the capsule and then contact the ends of opposite fibers to form Y sutures in the visual axis of the lens^6–8^. Each generation of secondary fibers forms a “growth shell” as successive shells are added during late development. Interior elongated fiber cells undergo a maturation process to eliminate intracellular organelles to form organelle-free mature fibers, which minimize light scattering along the visual axis of the lens.

The development, structure, and function of lens sutures are integrated into the lens’s ability to remain transparent, focus light and adjust optical power throughout life. Lens sutures, formed by interconnected network and microfilaments to stabilize fiber ends^9^, maintain the lens’ shape during accommodation, influence optical properties, and were conventionally thought to provide pathways for fluid flow and nutrient delivery as a part of lens microcirculation. Disrupted lens fiber cell organization and abnormal suture development are associated with pathological light scattering conditions such as cataracts^7^. Lens cellular abnormalities and molecular pathological properties have been primarily studied using dissected lenses *in vitro*. Therefore, it is necessary to apply high-resolution imaging technology to directly visualize *in vivo* native cellular and subcellular pathological information of lens cells and structures in live states.

Optical microscopy, owing to its subcellular resolution and non-invasiveness, has become a powerful tool for studying living organisms. Among conventional imaging modalities, 2-photon fluorescence microscopy (2PFM)^10^ possesses unique advantages for studying the mouse eye^11–13^, including the lens^13^. The nonlinear absorption process imparts intrinsic optical sectioning capabilities to 2PFM, allowing for depth-resolved 3D imaging of the mouse eye lens. Moreover, utilizing near-infrared (NIR) excitation that scatters less, 2PFM is well-suited for studying normal and pathological lenses. One challenge faced by *in vivo* 2PFM mouse eye imaging is optical aberrations introduced by the imperfect mouse eye, resulting in degraded resolution and contrast. Incorporating adaptive optics (AO), we recently demonstrated the first 2PFM imaging of the mouse eye lens *in vivo* and discovered unreported lens features^13^. However, this AO-2PFM system had a limited imaging field of view (FOV) and imaging speed, which did not fully capture the organization of lens fiber cells nor provide significant insights into the diseased mouse model.

In this study, utilizing a 2PFM system with resonant scanners to achieve higher imaging speed over a larger FOV, we have examined the structure and function of lens sutures between transparent (WT) and cataractous (KLPH-KO) lenses. KLPH, encoded by the LCTL (γ-Klotho) gene, is a member of the Klotho family (with KL/α Klotho and KLB/β Klotho) and a type I membrane glycoprotein. Mouse klotho gene mutations lead to aging syndrome^14^. Lens KLPH mRNA and protein are highly expressed in the equatorial epithelium and elongating fibers^15,16^. Confocal images of KLPH-KO lenses *in vitro* showed loose or open double-Y- or X-shaped lens sutures and eventually developed age-related cortical cataract^15^. Although it is unclear how KLPH deficiency leads to loosened sutures, KLPH-KO mice seem to be a valuable model to test the hypothesis for the role of lens sutures in lens microcirculation and optics^17^.

The results of these *in vivo* imaging experiments of live mice reveal various lens suture pathological features that were underappreciated with previous imaging methods: including reduced light transmission efficiency, disorganized suture lines, larger and multiple voids at the suture junction, more side voids in addition to central voids, and disorganized fiber cells and their degradation. The imaging data reveal that the central void area, which was observed but whose causes were unknown in our previous study^13^, is composed of mostly healthy (i.e., membrane integrity undisrupted) cells, along with degrading cells and extracellular space. To our knowledge, this work represents the first *in vivo* 2PFM investigation of fine pathological features in the lens sutures and provides new morphological markers for evaluating lens optical pathology and suture cataracts. We have further examined whether the diffusion pathway of FITC-Dextran is present along the lens anterior suture region using *ex vivo* imaging of FITC-Dextran dye-incubated lenses.

## Results

### 2PFM imaging of fiber cell organization and its changes in the KLPH-KO lens

Lens sutures typically exhibit characteristic “Y” or “double Y” patterns in mice^18–20^. Three-dimensional imaging of the lens suture structures has been challenging due to a lack of optical sectioning capability in conventional widefield microscopy. Although confocal fluorescence microscopy offers optical sectioning capability, its application has been restricted to dissected, thin lens sections due to limited penetrating ability, except when moving to longer wavelengths^21^. Since a previous study reported that KLPH-KO lenses had a loose suture defect based on the *in vitro* images^13^, we thought to use a 2PFM system with resonant scanners to achieve higher imaging speed over a larger FOV to examine the 3D suture structures in normal and loose-suture lenses. In addition, we wanted to evaluate if this *in vivo* lens imaging technology would allow us to test a long-standing hypothesis that the lens suture provides an effective diffusion pathway for maintaining lens homeostasis in the lens core^22,23^.

By utilizing 2PFM, we conducted *in vivo* imaging of age-matched mouse lenses over large volumes (approximately 800 × 800 × 800 µm^3^), revealing that suture patterns vary at different depths along the optical axis in both WT and KLPH-KO mice, as also seen in their depth-encoded projections. Consistent with previously published papers^18,19^, WT lens sutures exhibited either “Y” or “double Y” patterns at different depths, roughly aligning with their “double Y” suture “envelopes” (**Fig. 1A, Fig. 1C, Fig. S1, and Fig. S3**).

**Fig. 1.**
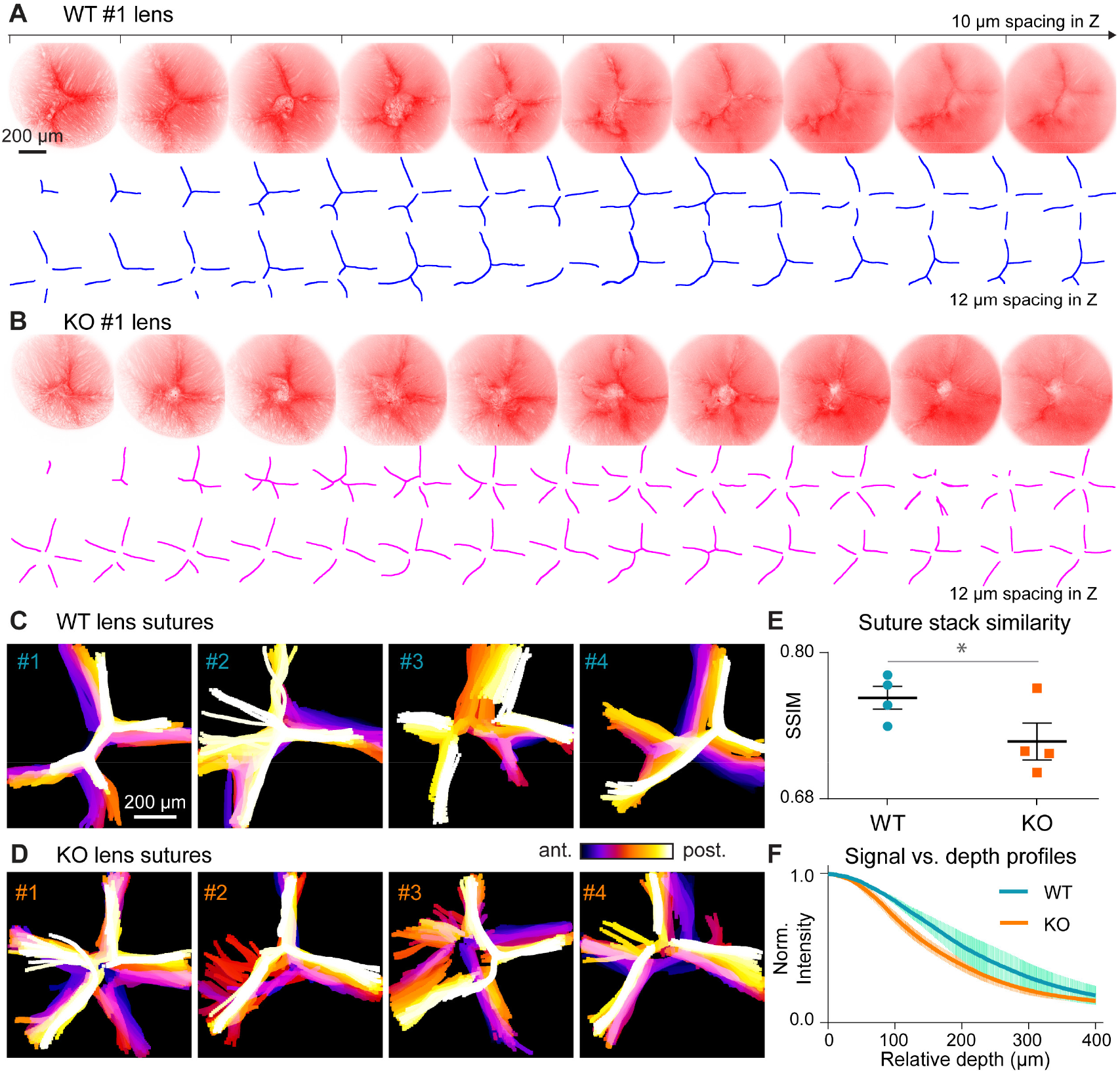
Organization KLPH-KO. **(A, B)** Representative 2PFM images of sutures (top) and manually extracted suture lines (bottom) with 10/12 μm depth interval, from a (**A**) WT and a (**B**) KLPH-KO mouse, respectively. Imaging field-of-view is 796.36 μm × 796.36 μm. **(C, D)** Depth-encoded projections of manually outlined suture patterns from a WT and KLPH-KO mice, respectively. **(E)** Quantification of suture pattern variations, expressed as the mean of structural similarity index value of images at each depth, using the mean projection as a reference. Error bars: ± std. The p value was determined using an unpaired one-sided Student’s t-test. **(F)** Normalized signal levels from WT and KLPH-KO mice lenses as a function of depth. Signal levels were normalized to the illumination power and the applied PMT gain. Shaded areas: std of signal measurements.

In contrast, KLPH-KO mice displayed diverse suture patterns including “Y”, “double Y”, and “star”, with variations appearing more randomly at different depths and misaligned suture planes (**Fig. 1B, Fig. 1D, Fig. S2, and Fig. S4**). The variability in suture patterns was quantified using the mean of image stacks’ similarity indices (SSIM) to the mean projection of the stack, showing a significant increase (p<0.05) in pattern randomization for KLPH-KO lenses (**Fig. 1E**).

This increased complexity and randomness in the suture structures of KLPH-KO mice could lead to reduced light transmission through the lens. Indeed, the fluorescence signal from KLPH-KO lenses decreased more rapidly than that from WT lenses, suggesting compromised lens transparency and potentially impacting vision quality (**Fig. 1F**).

### Characterization of voids and vacuoles along the lens suture in FITC-Dextran incubated lenses *in vitro*

Our previous work^13^, applying 2PFM for *in vivo* imaging of mouse lenses, identified large voids and enlarged vacuoles in the anterior part of the KLPH-KO mouse lenses. High-resolution 2PFM further confirmed the presence of these voids and vacuoles, but whether they are intra-cellular or extra-cellular structures remains undetermined.

By imaging both WT and KO mice, we observed that both lenses exhibit such voids where sutures meet, as well as enlarged vacuoles throughout the lens volume *in vivo* (**Fig. 2A**). To investigate the physiological nature of these structures at the lens suture and their potential involvement with intralenticular diffusion pathways, we incubated enucleated fresh WT and KO lenses with a FITC-Dextran solution overnight before 2PFM imaging. The results show that the green dye successfully penetrated the lens, reaching a depth of over 300 µm (**Fig. 2B**). Since FITC-Dextran (MW 2M/10K) is impermeable to the lens cell plasma membrane, which is labeled with tdTomato in both WT and KO mice, diffused FITC-dextran signals detected by two-color 2PFM images, were mainly localized in extracellular spaces, with a few exceptions where some lens epithelial and fiber cells might be damaged to lead to positive FITC-Dextran inside. We speculated such cell damage might be associated with the long incubation of fresh lenses with dyes *in vitro*. However, it is unknown how some FITC-Dextran got into those cells **(Fig. 2C**).

**Fig. 2.**
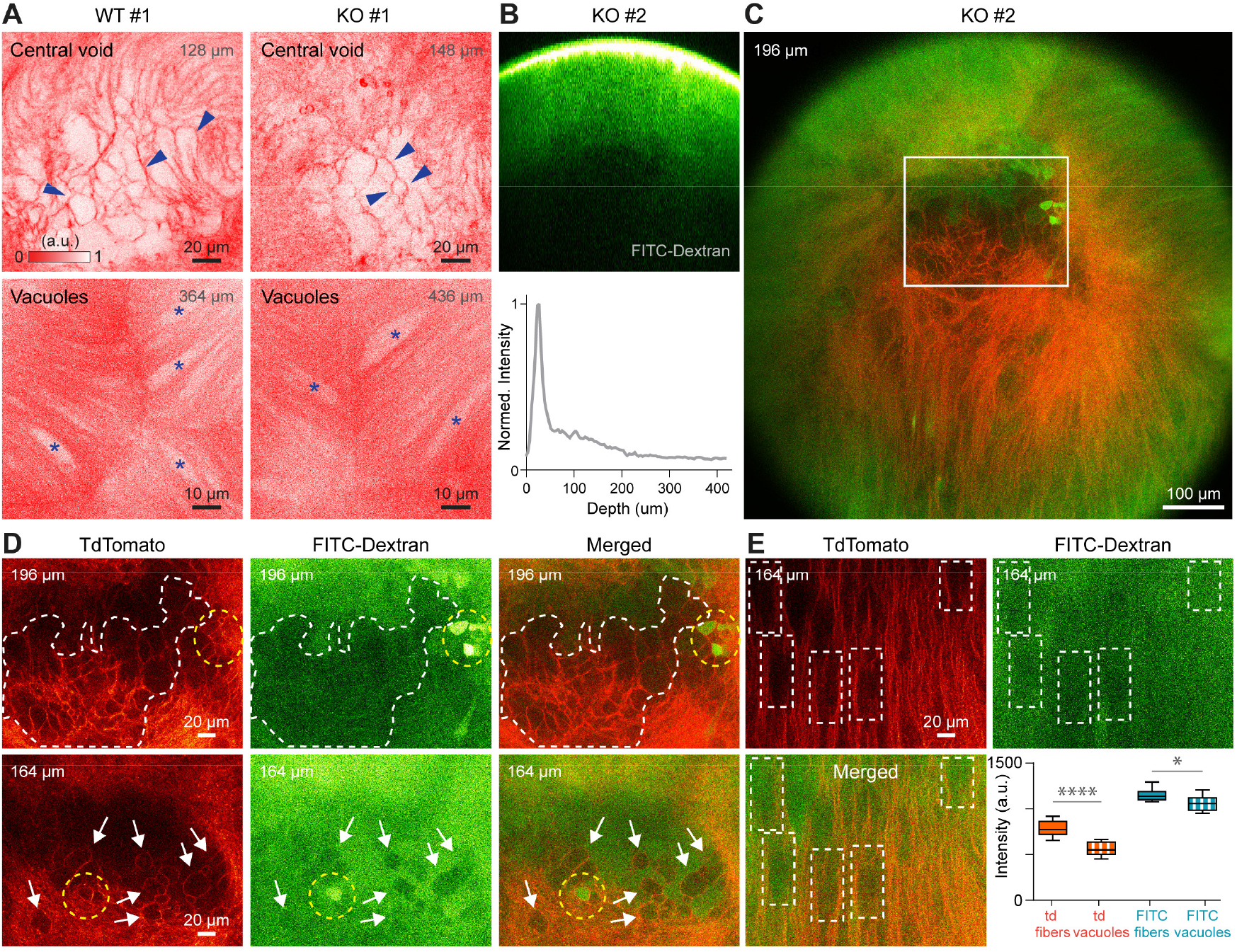
2PFM imaging of central voids and vacuoles in the mouse lens. **(A)** *In vivo* single-plane 2PFM images of the central region at the conjunction of suture lines acquired from a WT (left) and a KLPH-KO (right) mouse lens, respectively. Central voids (blue arrow heads) and vacuoles (asterisk) are highlighted. **(B)** Top: cross-sectional view of a FITC-Dextran immersed KO lens. Bottom: green fluorescence signal measured by 2PFM. **(C)** Merged 2-color 2PFM image of a tdTomato-labeled KO lens incubated with FITC-Dextran. **(D)** Single-plane 2PFM images of the central region at the conjunction of suture lines in the (left) red and (middle) green channels as well as (right) the merged image, acquired at (upper) 196 µm and (lower) 164 µm, respectively. **(E)** Single-plane 2PFM images of the vacuoles acquired in both red and green channels as well as the merged image. Bottom right: signal comparison using the mean intensity calculated from 9 fiber and 9 vacuole regions measured in both channels. All images were adjusted for better visualization. [Representative data from three animals.]

At a depth of 196 µm from the lens surface, we observed that the central void mostly belongs to an area with a globally decreased green signal (**Fig. 2D**, upper middle, white dashed shape). We speculated that “voids” in this region, defined by tdTomato-labeled plasma membranes (**Fig. 2D**, upper left, white dashed shape) and minimal FITC-Dextran labeling areas, were likely intracellular regions of tightly tethered fiber cell bundles with impermeable extracellular spaces. Notably, there were a few cells at the same depth near this region labeled by intense FITC-Dextran green signals (**Fig. 2D**, upper row, yellow circles). These FITC-Dextran positive cells likely resulted from cells with limited damage during incubation *in vitro*. Similar observations were made at another depth (164 µm), where some cells had lower signals (**Fig. 2D**, lower row, white arrows) and one cell was labeled with FITC-Dextran (**Fig. 2D**, lower row, yellow circle). These findings suggest that the central region, or central voids, formed at the conjunction of the suture lines, mainly comprises tightly apposed and irregularly shaped ends of lens fiber cells, with only very limited fiber cells (likely damaged or unhealthy) containing positive FITC-Dextran dyes (**Fig. 2 and Fig. S5**).

Two-color 2PFM imaging data also revealed that the enlarged “vacuoles” contained reduced green signals compared to adjacent lens fibers (**Fig. 2E**, 5 dashed white boxes). The mean intensity of fluorescence signals calculated from 9 fiber and 9 vacuole regions confirmed that these “vacuoles” had reduced green FITC-Dextran signals and decreased tdTomato signals (**Fig. 2E**, lower right). Since normal mouse lenses must be composed of elongated fibers with minimal extracellular spaces^4,24–27^, lens fibers also form these “vacuoles”. Therefore, we speculate that these “vacuoles” likely resulted from lens fiber cell regions containing less membrane-tagged tdTomato that are tightly opposed to each other to reduce the extracellular permeability of FITC-Dextran.

### Pathological features in the KLPH-KO lenses

Lens sutures are formed by the contacts of the ends of elongated fiber cells. Disrupted contact stability and/or altered morphological and/or biophysical properties at the ends of elongated fiber cells are likely to cause suture cataract formation in KLPH-KO lenses. We have then focused on *in vivo* imaging of this suture region to identify pathological evidence in KLPH-KO in comparison with WT lens control (**Fig. 3**).

**Fig. 3.**
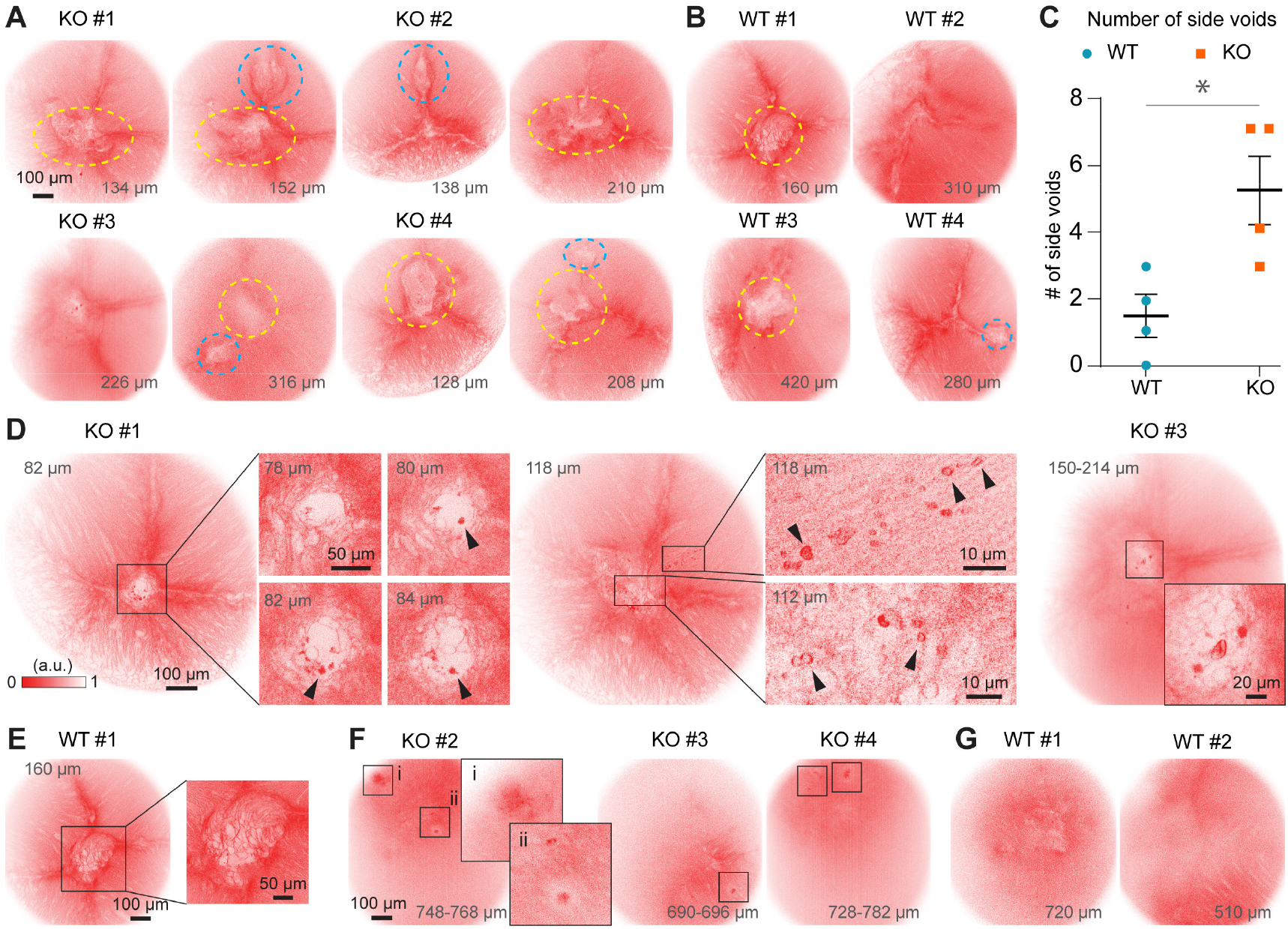
2PFM imaging of four KLPH-KO and WT lenses. **(A)** Four KO lenses showing central and side voids formed at the central conjunction of suture lines (yellow circles) and between two suture lines (cyan circles), respectively. Fewer side voids are seen in four WT mice **(B)** and are confirmed by an unpaired one-sided Student’s t-test **(C). (D, E)** Representative lens images from two KO mice showing degraded fiber cells (black arrowheads), whereas this is not observed in the WT mouse. **(F, G)** Representative lens images showing degraded fiber cells in KO but not in WT at deeper depth.

Comparing four KLPH-KO (**Fig. 3** and **Fig. S2**) with four WT (**Fig. 3, Fig. S1 and Fig. S6**) lenses, we observed that KLPH-KO lenses had at least one more void within the imaging volume (**Fig. 3A**, cyan dashed circles), apart from the central one (**Fig. 3A**, yellow dashed circles) formed at the conjunction of suture lines, while fewer additional voids were observed in WT lenses (**Fig. 3B and Fig. 3C**). These additional voids might further impair light transmission and evolve into areas with additional pathological fiber cells. In addition, we observed that KLPH-KO lenses have larger central voids that tend to exhibit irregular shapes. Within these voids, a remarkable pathological feature was identified in the KLPH-KO lenses: the formation of 2~5-µm-diameter ring-like or amorphous structures (**Fig. 3D**, insets, black arrowheads). Given membrane-tethered tdTomato proteins, we speculate these structures are membrane protein remnants/aggregates from pathological fiber cells. Moreover, these features were found both within and outside the central voids’ region (**Fig. 3D**, insets at 118 µm depth in mouse KO #1). In contrast, no such feature was observed in the four WT lenses imaged (**Fig. 3E, Fig. S1, and Fig. S6**). Furthermore, in deeper regions of the KLPH-KO mouse lenses, small areas with concentrated fluorescence expression were observed (**Fig. 3F**), a feature not seen in the WT lenses (**Fig. 3G and Fig. S6**).

## Conclusion and Discussion

The intrinsic optical sectioning capability of 2PFM allowed us to observe depth-varying suture patterns in both WT and KLPH-KO mouse lenses. Based on precise graphic analysis and statistical data of the suture pattern and organization of 2PFM images between WT and KLPH-KO lenses, we found that the suture patterns in KLPH-KO mouse lenses exhibit greater diversity, increased complexity, and irregular randomness, along with pathological remnants of fiber cell ends, all of which likely contribute to reduced light transmission and promote lens opacity. This work suggests that KLPH plays a critical role in the regulation of lens suture formation and disrupted sutures likely contribute to cataract formation in KLPH-KO lenses. In addition, a well-established suture pattern is important for light transmission of the lenses.

Through *ex vivo* 2PFM imaging of dye-incubated lenses, we confirmed that the central voids comprise a mixture of fiber cells that are tightly apposed, have irregularly shaped ends, and contain tighter extracellular spaces. The 2PFM results of lens FITC-Dextran dye incubation indicate that the opposite ends of fiber cells tightly contact at the suture, and no obvious diffusion pathway for high molecular weight dextran dye is present between extracellular spaces along lens suture lines in both WT and KLPH-KO lenses. Thus, the extracellular spaces of lens peripheral elongating and elongated fibers mediate the dye diffusion of over 300 µm into the lens (**Fig. 2B**). We did not observe an open suture in KLPH-KO lenses *in vivo* and *in vitro*, unlike the previous paper, which was based on the imaging analysis of fixed lenses^28^. Tdtomato remnants and aggregates in high-resolution 2PFM images indicate pathological conditions of fiber cell ends along suture regions in KLPH-KO. This work further demonstrates that KLPH is important for maintaining the integration of fiber cell ends and suture formation. Additionally, pathological changes detected by 2PFM images are new and valuable morphological markers for cataract formation *in vivo*.

We have also observed extensively enlarged “vacuoles” in live lenses at different depths that seem to have unique structures formed by a bundle of fibers (consisting of 4-5 fibers according to their 10-20 um in diameter) that have reduced tdTomato labeling and tighter extracellular spaces. These “vacuoles” seem unlikely to be caused by enlarged fiber size in selective regions of these elongated fibers or by the plasma membrane fusion of attached fibers at specific regions^18,29^. Given various pathological features in *in vivo* images of KLPH-KO mouse lenses, including larger, more irregular, and multiple central voids containing membrane-enclosed remnants of pathological fibers, indicating a disruption in the organization of lens fiber cells, we hypothesize that KLPH plays an important role in supporting the overall integrity and stability of elongated fiber cells, besides the regulation of fiber cell ends at the sutures. At the current stage, the molecular mechanisms for how KLPH protein supports the formation and/or stability of lens suture patterns remain unknown.

In conclusion, this study not only demonstrates the utility of 2PFM in revealing the complex 3D organization of lens fiber cells *in vivo* but also provides valuable insights into the pathological changes that occur in pathological lenses. The identification of specific structural abnormalities in KLPH-KO mice offers a new perspective on the mechanisms underlying lens transparency and cataract formation, laying the groundwork for future research aimed at preventing or reversing these conditions *in vivo*. Furthermore, our work highlights the potential of advanced optical imaging techniques in contributing to a deeper understanding of eye lens biology and pathophysiology, offering promising avenues for therapeutic intervention in lens-related vision impairments.

## Methods and Materials

### Animal Use

All animal experiments were conducted according to the National Institutes of Health guidelines for animal research. Procedures and protocols (AUP-2020-06-13343) were approved by the Institutional Animal Care and Use Committee at the University of California, Berkeley.

### Mouse models

The membrane-targeted tdTomato-expressing (007676, ROSA^mT/mG^) mice were purchased from the Jackson Laboratory. The klotho-related protein KLPH (lctl) knockout (KO) mice were a gift from Dr. Melinda Duncan at the University of Delaware. The KLPH-KO mice were originally generated by Dr. Graeme Wistow at the National Eye Institute. In this study, we used the membrane-targeted tdTomato-expressing mice as the wildtype (WT) control mice^30^, and we used the backcrossed tdTomato-expressing and KLPH-KO mice as the diseased mice^28^. All *in vivo* imaging experiments were performed on mice at the age of 7 or 8 months. The genotype, sex, and age at imaging are summarized in Supplementary **Table S1**.

### 2-photon fluorescence microscope (2PFM)

For *in vivo* mouse lens imaging, we utilized a commercial 2PFM microscope (Thorlabs Bergamo^®^ II multiphoton). A femtosecond Ti:Sapphire laser (Coherent, Chameleon Ultra II) was used as the excitation source for all experiments. The laser was tuned to 1000 nm for *in vivo* imaging of the tdTomato-expressing mouse eye lenses, and it was tuned to 920 nm for *ex vivo* imaging of the FIFC-Dextran-incubated tdTomato-expressing mouse eye lenses. A 25× water-dipping objective lens (Olympus, 1.05 NA, 2 mm WD) was used to focus the excitation light into the mouse eye lens and collect the emitted fluorescence for imaging. Data acquisition and hardware were controlled using the ThorImage software. The imaging experiment settings are summarized in Supplementary **Table S1**.

### *In vivo* mouse eye lens imaging

*In vivo* imaging experiments were carried out on mice under light isoflurane anesthesia (0.5-1% by volume in O_2_). Prior to imaging, the mouse was stabilized on a bite-bar on a custom-made stage with a dual-axis goniometer stage (Thorlabs, GNL 18) and was aligned to make the pupil face up. The mouse pupil was dilated with 2.5% phenylephrine hydrochloride (one drop, Paragon BioTeck, Inc.) and 1% tropicamide (one drop, Akorn, Inc.). Following that, to prevent cornea drying and clouding, eye gel (Genteal) was applied between the eye and a cover glass (Fisherbrand^®^, No. 1.5, 0.16-0.19 mm thick). The cover glass was mounted on a holder to minimize the effect of eye motion and was carefully aligned to be perpendicular to the excitation light. During imaging, the mouse’s body temperature was maintained using a hand warmer. The correction collar of the objective lens was adjusted to minimize spherical aberration induced by the cover glass.

### *Ex vivo* mouse eye lens imaging

Eyes were retrieved and dissected in culture media made of M199 (Invitrogen, Carlsbad, CA) supplemented with Hepes (Invitrogen, Carlsbad, CA) at 37 degrees Celsius to extract the lens. Dissected lenses were then incubated in the culture media with either 2,000,000 MW FITC-Dextran (Sigma-Aldrich, St. Louis, MO) or 10,000 MW FITC-Dextran (Molecular Probes, Eugene, OR) at different concentrations. The lenses were incubated at 37 degrees Celsius, 5% CO_2_, and 95% humidity for 20 or 24 hours (Supplementary **Table S1**). Dissected lenses were then washed with culture media 3 times before imaging. During imaging, lenses were placed in wells with the M199 culture media as the immersion medium.

### Image processing and analysis

All image processing, visualization, and analysis were performed in ImageJ (NIH, Bethesda, MD, USA)^31^ and MATLAB (Mathworks, Natick, MA, USA). Images of suture lines were hand-drawn at different depths in ImageJ.

## Acknowledgements

We thank Dr. Santosh Paidi, Dr. Yuxing Li and Dr. Iksung Kang for helpful discussions. This work was supported by US National Institutes of Health grants U01NS118300 and U01NS137449 (Q.Z., J.Z., and N.J.), NIH/EY013849 (T.P., C.H.X., and X.G.).

## Author contributions

X.G. and N.J. supervised this project. Q.Z. and J.Z. conducted *in vivo* mouse eye lens imaging experiments. Q.Z., J.Z., and T.P. conducted *ex vivo* lens imaging experiments. T.P. contributed to the sample preparations.

Q.Z. and J.Z. analyzed imaging results. C.X. provided mouse lines and mentoring support. Q.Z., J.Z., T.P., and X.G. wrote the manuscript with input from all authors.

## Data and material availability

All data generated or analyzed in this study are included in the manuscript and supplementary materials. Source data may be obtained from the authors upon reasonable request.

## Supplementary Materials

**Figure S2.**
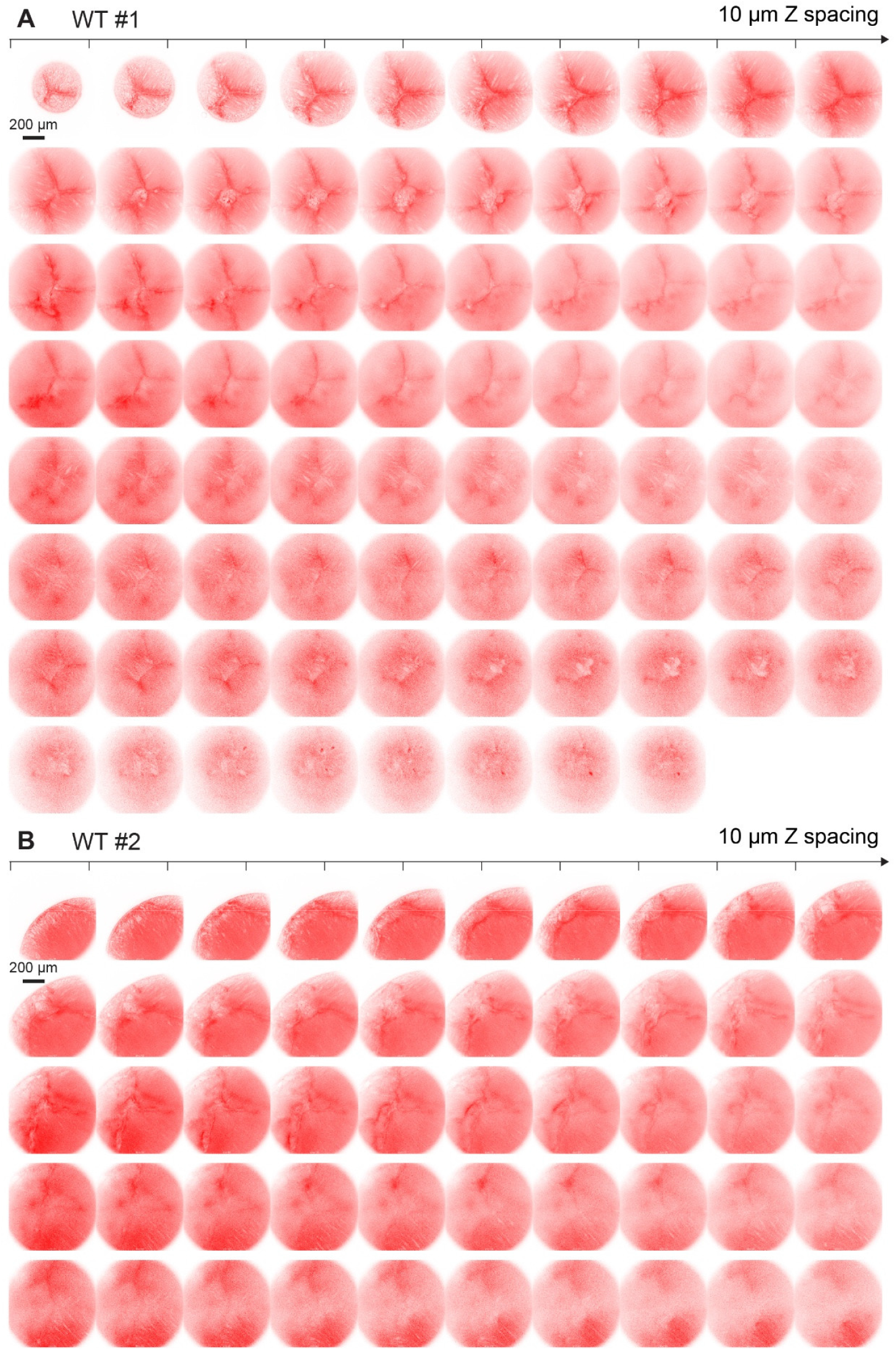
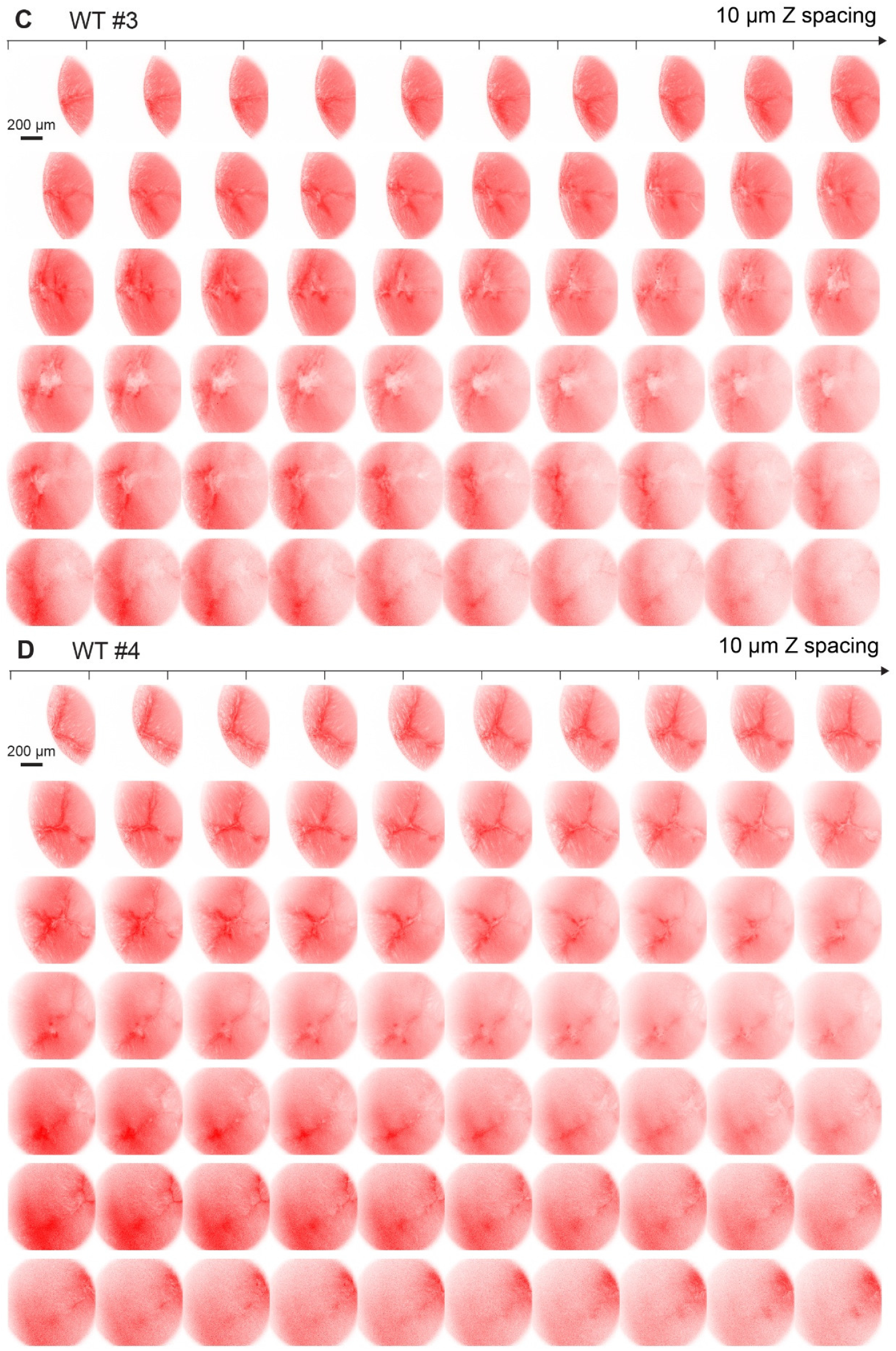
2PFM lens images of the KLPH-KO mice acquired at different depths.

**Figure S2.**
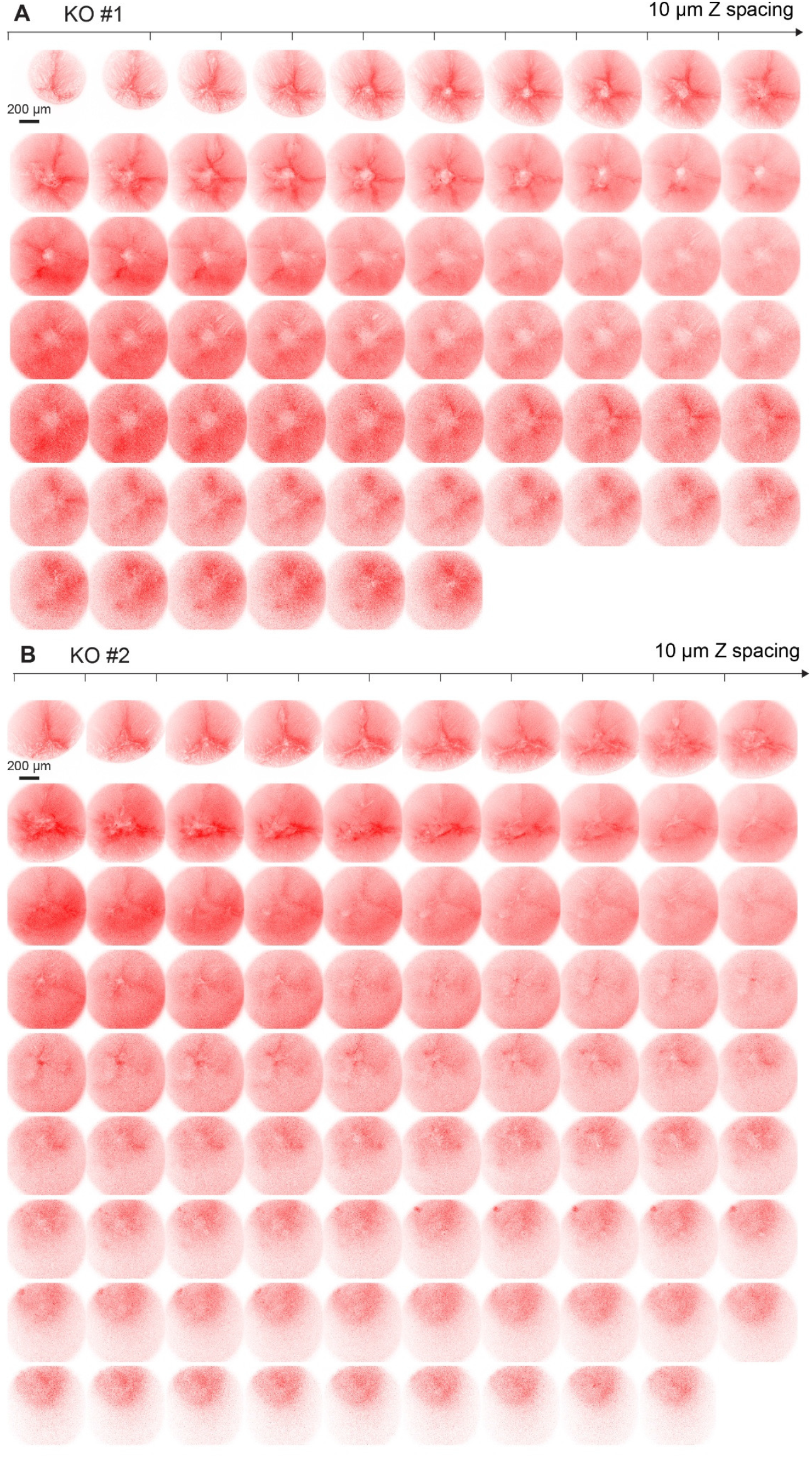
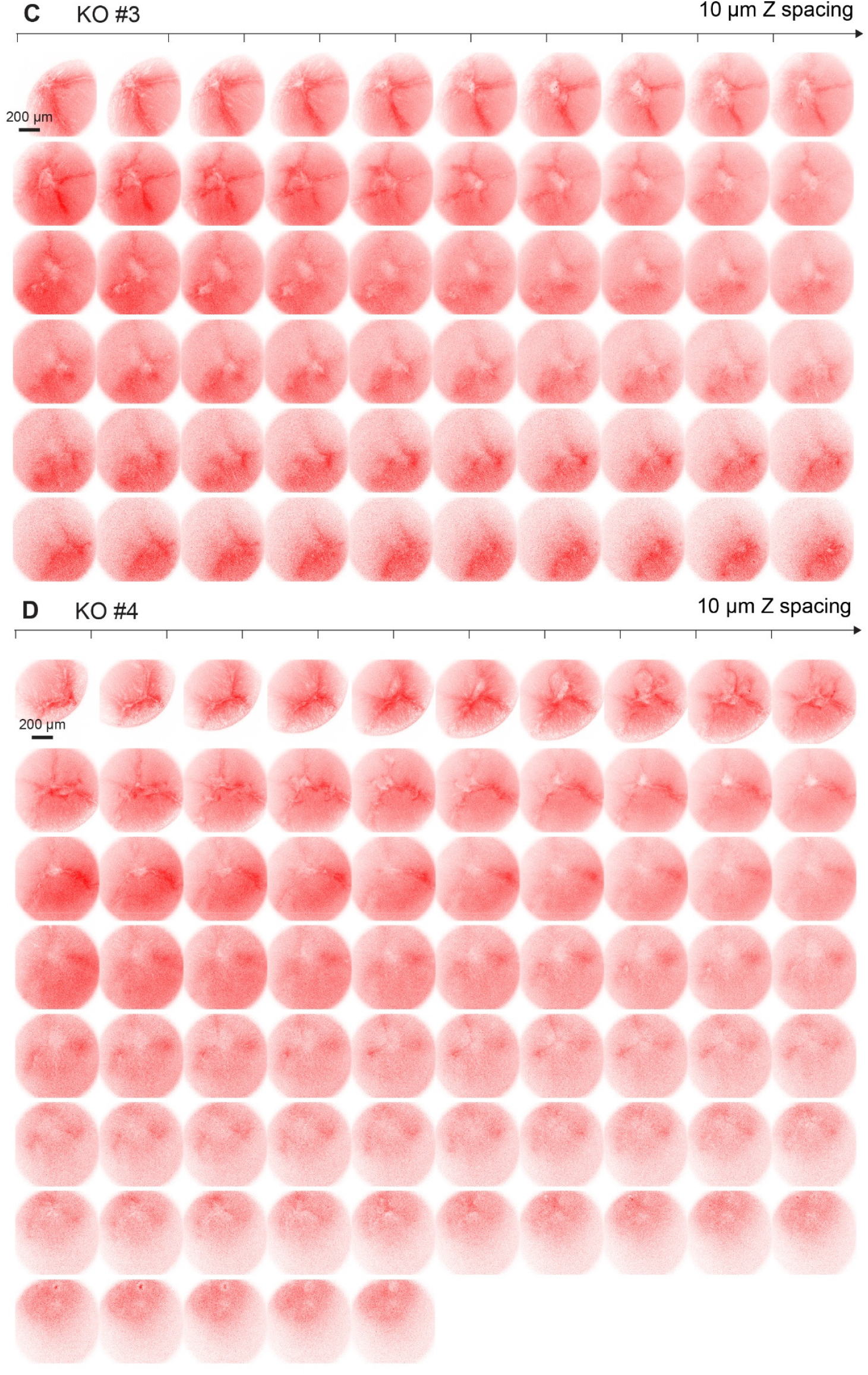
2PFM lens images of the KLPH-KO mice acquired at different depths.

**Figure S3.**
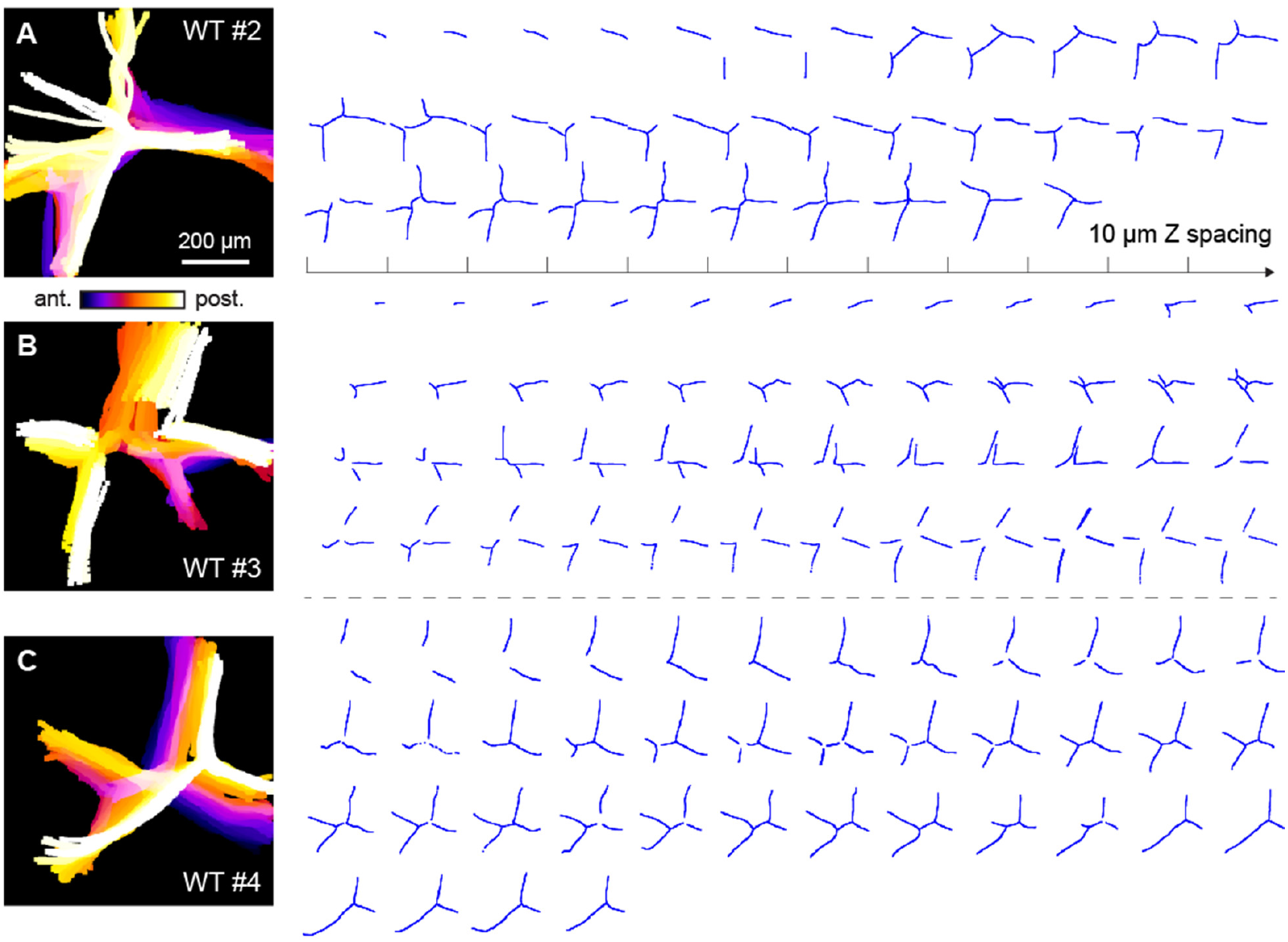
Suture lines of the wildtype (WT) mice at different depths. **(A-C)** Left: Depth-encoded projections of hand-drawn suture stacks from three other WT mice. Right: Suture lines at various depth at 10 µm spacing in Z. Scalebar represents 200 µm and applies to all the images.

**Figure S4.**
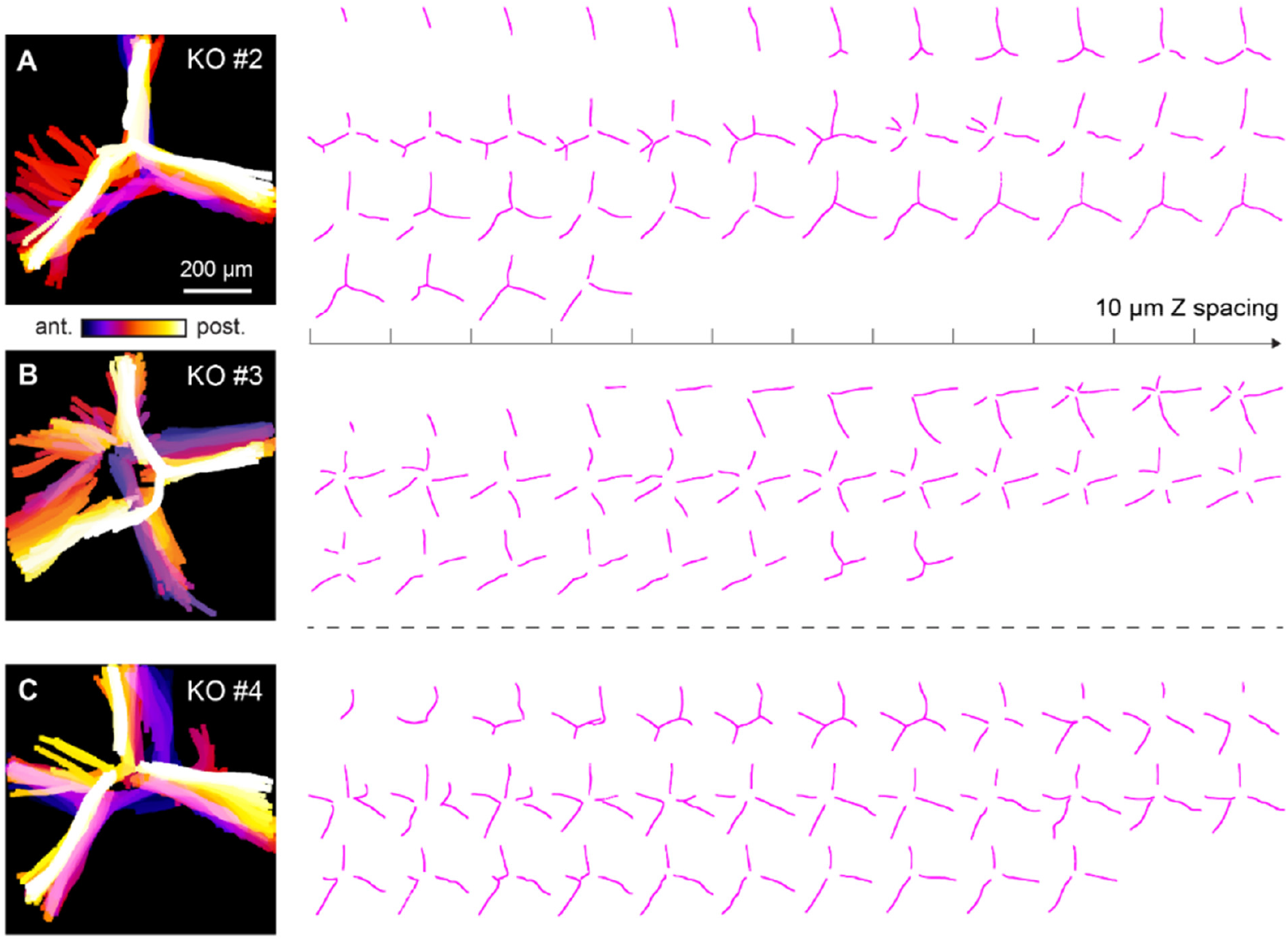
Suture lines of the KLPH-KO mice at different depths. **(A-C)** Left: Depth-encoded projections of hand-drawn suture stacks from three other KO mice. Right: Suture lines at various depth at 10 µm spacing in Z. Scalebar represents 200 µm and applies to all the images.

**Figure S5.**
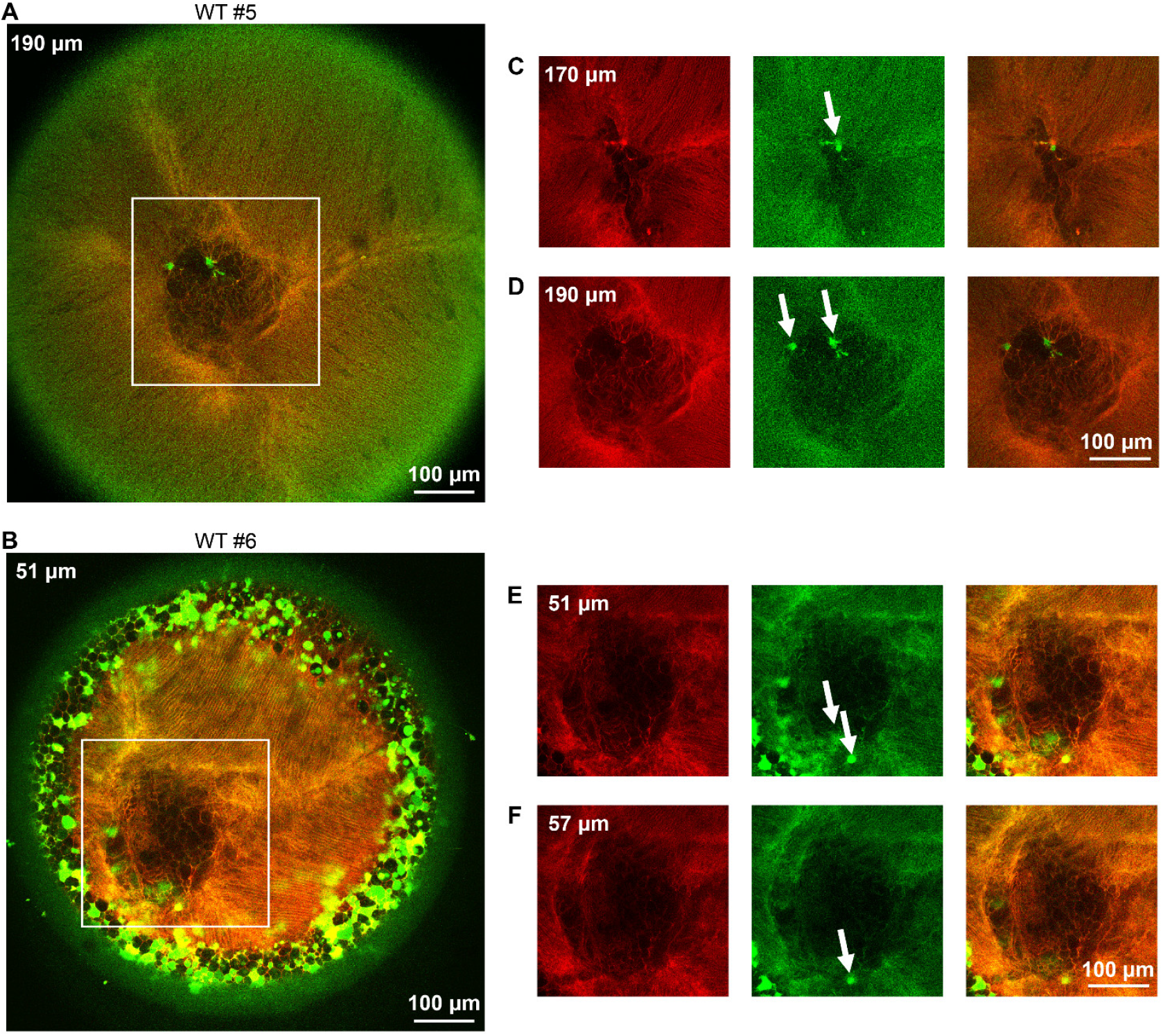
2PFM imaging of central voids in WT #5 and WT #6 mouse lenses. (A-B) Merged 2-color 2PFM images of WT lenses incubated with FITC-Dextran. (C-F) Single-plane 2PFM images of the lens suture conjunction region (white boxes in A and B) at different depths. White arrows denote FITC signal in the green channel but not in the red channel, indicating dye penetration ability but without entering the suture conjunction area. Left: TdTomato imaged in the red channel. Middle: FITC-Dextran imaged in the green channel. Right: Merged images. Scalebars 100 µm.

**Figure S6.**
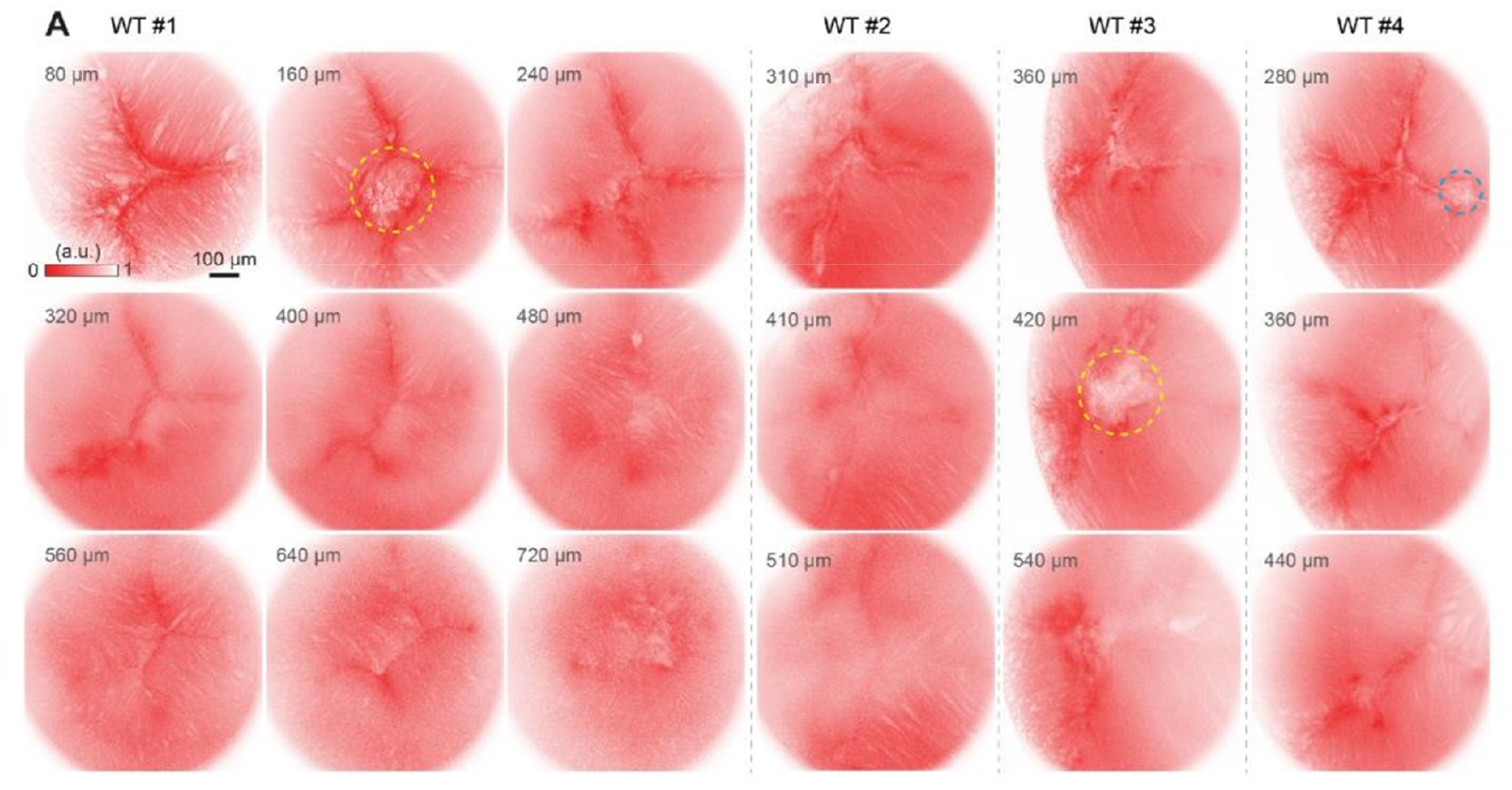
2PFM imaging of four wildtype (WT) lenses at different depths with void structure highlighted. Yellow circles: central void at the conjunction of suture lines. Blue circle: lens void formed between two suture lines. Scalebar represents 100 µm and applies to all the images.

**Table S1.** Animal information and experimental settings for all experiments.

|  |  | Genotype | Sex | Age at imaging | Imaging parameters |
| --- | --- | --- | --- | --- | --- |
| *In vivo*<br>imaging | WT #1 | tdTomato × WT | Male | 7 months and 11 days | FOV: 796.36 μm × 796.36 μm<br>Depth range: 600-980 μm<br>Z step size: 2 μm<br>Frames averaged at each depth: 15<br>Imaging wavelength: 1000 nm |
|  | WT #2 | tdTomato × WT | Male | 7 months and 11 days |  |
|  | WT #3 | tdTomato × WT | Male | 7 months and 11 days |  |
|  | WT #4 | tdTomato × WT | Male | 7 months and 11 days |  |
|  | KO #1 | L3: LCTL (-/-) × LCTL (-/-) | Female | 8 months and 2 days |  |
|  | KO #2 | L3: LCTL (-/-) × LCTL (-/-) | Female | 8 months and 2 days |  |
|  | KO #3 | L3: LCTL (-/-) × LCTL (-/-) | Female | 8 months and 2 days |  |
|  | KO #4 | L3: LCTL (-/-) × LCTL (-/-) | Male | 8 months and 3 days |  |
| *Ex vivo*<br>imaging | KO #2 | L3: LCTL (-/-) × LCTL (-/-) | Female | 13 months and 2 days | FOV: 796.36 μm × 796.36 μm<br>Depth range: 420 μm<br>Z step size: 4 μm<br>Frames averaged at each depth: 20<br>Imaging wavelength: 920 nm<br>Dye used: 2,000,000 MW FITC-Dextran<br>Incubation time: 24 h |
|  | WT #5 | tdTomato × WT | Male | 7 months and 29 days | FOV: 796.36 μm × 796.36 μm<br>Depth range: 520 μm<br>Z step size: 5 μm<br>Frames averaged at each depth: 20<br>Imaging wavelength: 920 nm<br>Dye used: 10,000 MW FITC-Dextran<br>Incubation time: 20 h |
|  | WT #6 | tdTomato × WT | Female | 3 months and 1 days | FOV: 796.36 μm × 796.36 μm<br>Depth range: 261 μm<br>Z step size: 3 μm<br>Frames averaged at each depth: 20<br>Imaging wavelength: 920 nm<br>Dye used: 10,000 MW FITC-Dextran<br>Incubation time: 20 h |

